# Partitioned Learning of Deep Boltzmann Machines for SNP Data

**DOI:** 10.1101/095638

**Authors:** Moritz Hess, Stefan Lenz, Tamara J Blätte, Lars Bullinger, Harald Binder

**Affiliations:** Institute of Medical Biostatistics, Epidemiology and Informatics (IMBEI), University Medical Center, Mainz, 55131, Germany; Department of Internal Medicine III, University Hospital of Ulm, Albert-Einstein-Allee 23, 89081 Ulm, Germany.

## Abstract

Learning the joint distributions of measurements, and in particular identification of an appropriate low-dimensional manifold, has been found to be a powerful ingredient of deep leaning approaches. Yet, such approaches have hardly been applied to single nucleotide polymorphism (SNP) data, probably due to the high number of features typically exceeding the number of studied individuals. After a brief overview of how deep Boltzmann machines (DBMs), a deep learning approach, can be adapted to SNP data in principle, we specifically present a way to alleviate the dimensionality problem by partitioned learning. We propose a sparse regression approach to coarsely screen the joint distribution of SNPs, followed by training several DBMs on SNP partitions that were identified by the screening. Aggregate features representing SNP patterns and the corresponding SNPs are extracted from the DBMs by a combination of statistical tests and sparse regression. In simulated case-control data, we show how this can uncover complex SNP patterns and augment results from univariate approaches, while maintaining type 1 error control. Time-to-event endpoints are considered in an application with acute myeloid lymphoma patients, where SNP patterns are modeled after a pre-screening based on gene expression data. The proposed approach identified three SNPs that seem to jointly influence survival in a validation data set. This indicates the added value of jointly investigating SNPs compared to standard univariate analyses and makes partitioned learning of DBMs an interesting complementary approach when analyzing SNP data.

## 1. Introduction

Identification of complex patterns comprising several single nucleotide polymorphisms (SNPs) is considered to be the key for better explaining phenotypic variability (Wei *et al.*, 2014). Applications with SNP data, such as genome-wide association studies (GWAS) or clinical cohorts, might thus benefit from identifying a low dimensional manifold providing a compact description of observed individuals (Bengio *et al.*, 2013).

Inherent to identifying a compact representation is learning of the joint distribution of the mostly high dimensional SNP data. The majority of all SNPs is bi-allelic which implies that a haploid SNP can be modeled as a Bernoulli variable. Consequently, the joint distribution of many SNPs could be expressed as a high-dimensional cross-table. While the joint distribution of a small number of Bernoulli variables could be estimated with a specific form of log-linear models, this approach is only feasible for up to about 20 variables.

In contrast, deep learning techniques allow to generate compact and accurate representations of high-dimensional binary data for a larger number of variables (Bengio *et al.*, 2013; Hinton and Salakhutdinov, 2006). They have been applied in a multitude of settings, with a limited number of applications also in bio-medical research (Quang *et al.*, 2014; Leung *et al.*, 2014; Angermueller *et al.*, 2016; Chen *et al.*, 2016). The general idea of deep learning is to employ a network structure for mapping observed variables into hidden variables by several network layers, each representing a non-linear transformation.

Deep Boltzmann machines (DBMs) (Salakhutdinov and Hinton, 2009) are a special type of Boltzmann machines, where observed variables and hidden variables in subsequent hidden layers are conditionally independent given the variables in the respective adjacent layers. Since deep Boltzmann machines impose restrictions on higher order interactions (Salakhutdinov and Hinton, 2009), they might provide an adequate stochastic model for a high-dimensional SNP cross-table. Still this leads to a large number of parameters even for a small number of variables, requiring sophisticated estimation techniques (Salakhutdinov and Hinton, 2012). As a consequence and similar to other deep learning techniques, the application of DBMs has so far been restricted to settings where the number of individuals is larger, and often much larger, than the number of observed variables (Chen *et al.*, 2014). General rules of thumb correspond to about *n* = 10 · *p* individuals required for *p* variables, naturally also depending on the specific network structure (Hinton, 2010).

In the following, we will investigate how DBMs could nevertheless be adapted for a large number of SNPs equal to or larger than the number of observed individuals. Specifically, we will consider settings with up to *n* · 5 variables. Such applications with a moderately large but not huge number of SNPs might e.g. be relevant when gene expression data has been used to already identify some potentially relevant genes, and the SNPs corresponding to these genes are to be investigated subsequently. One such application will be presented with data from clinical cohorts of acute myeloid leukemia (AML) patients (Hieke *et al.*, 2016a,b; The Cancer Genome Atlas Research Network, 2013).

To enable estimation of DBMs even for a rather large number of SNPs, we will propose an approach for identifying a partition into clusters of SNPs whose joint distribution should be learned in order to derive the overall joint distribution. Thereby we avoid the costly parameter estimation for practically independent SNPs. Specifically, we will use a regularized regression approach, stagewise regression (Tutz and Binder, 2007), for performing variable selection for each SNP as a response variable with respect to all other SNPs. The resulting variable selection forms the basis for clustering SNPs and for subsequently partitioning the parameter estimation of an overall DBM into one sub-DBM per cluster.

The resulting model for the joint distribution of SNPs could be useful for different tasks. While deep learning research has put a strong emphasis on prediction performance (Krizhevsky *et al.*, 2012; Ciregan *et al.*, 2012; Graves *et al.*, 2013), which in the present context would correspond to predicting disease states or risk of death based on high-dimensional SNP data, extraction of identified patterns from deep networks has received less attention. Yet, this is important for understanding the interplay of SNPs. Therefore, we will propose an approach for extracting SNP patterns from a DBM. Specifically, we will combine statistical testing on the hidden nodes with variable selection, provided by stagewise regression, to identify significant SNPs linked to a phenotype under type 1 error control.

In the Methods section, we will briefly review parameter estimation for DBMs, indicating some of our experience in tuning the approach for our application, before introducing the partitioning approach. Subsequently, we show how to extract significant SNPs from the resulting DBMs. A case-control simulation design is described in the subsequent section, and used to evaluate the performance of our proposals, in particular compared to standard univariate testing. In the application to SNP data from acute myeloid leukemia patients, the approach is illustrated for time-to-event endpoints. Finally, we will provide concluding remarks and indicate potential future extensions and generalizations.

## 2. Methods

In the following we consider settings where a phenotype is to be linked to a potentially large number of SNPs. For example, the case-control status takes the role of a binary phenotype in genome-wide association studies (GWAS), or survival might be the endpoint of interest in a clinical cohort. We assume dominant SNP effects, which simplifies the diploid SNP encoding to a binary variable, where 0 corresponds to no minor or risk allele while 1 corresponds to at least one minor or risk allele. The results can easily be generalized to other SNP codings, as will be discussed later.

### 2.1. Deep Boltzmann machines

The joint distribution of *p* single nucleotide polymorphisms (SNPs) could in principle be described by a log-linear model. Yet, such models are only suitable for estimating the joint distribution of a small number of variables. In contrast, deep Boltzmann machines (DBMs) (Salakhutdinov and Hinton, 2009) provide a model for the joint distribution of a large number of Bernoulli variables, also employing hidden nodes in a multi-layer network. In the following, we will restrict ourselves to networks with two hidden layers (with vectors **h**^(1)^ and **h**^(2)^ indicating the activation of these layers). Using this DBM we aim to model the probability of the visible nodes **v** corresponding to the *p* SNP variables, i.e. the underlying joint distribution. The DBM specifies the log-probability as

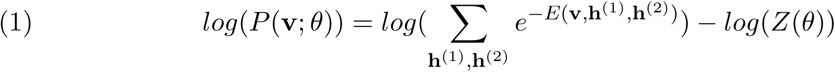

where *θ* corresponds to the parameters of the DBM. *E* is the energy function

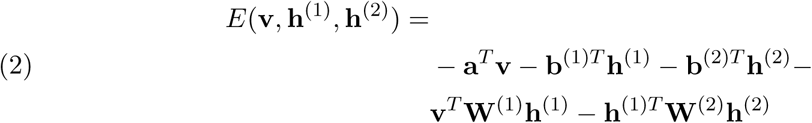

where **W**^(1)^ and **W**^(2)^ correspond to the weight matrices connecting **v** with **h**^(1)^ and **h**^(1)^ with **h**^(2)^ respectively. **a**, **b**^(1)^ and **b**^(2)^ are the so-called bias vectors, corresponding to intercept terms. The log-partition function

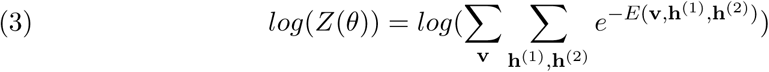

normalizes the probability.

With their layered architecture, DBMs are able to employ high-level information to constrain parameters that capture lower-level information in order to derive a better representation of the data (Salakhutdinov and Hinton, 2012).

For parameter estimation, we do as recommended in Salakhutdinov and Hinton (2009, 2012). Layer-wise pre-training is employed to get closer to a solution that optimizes the likelihood. For pre-training the deep Boltzmann machine is treated as two stacked restricted Boltzmann machines, RBM1 and RBM2, whose weight matrices **W**^(1)^, **W**^(2)^ are estimated consequently using contrastive divergence (Hinton, 2002). After the parameters of RBM1 are estimated, RBM2 is trained on the activations of **h**^(1)^ which are derived by passing the training data through RBM1. By adding more layers, the variational lower bound of the likelihood of the data is increased while the likelihood itself may not necessarily be increased.

Joint refinement is performed by a combination of mean field approximation of the data dependent distribution by variational learning (Neal and Hinton, 1998) and Gibbs sampling with parallel Gibbs chains for the approximation of the distribution defined by the DBM. The joint refinement finally improves the lower bound of the likelihood of the data (Salakhutdinov and Hinton, 2012).

The performance in modeling the joint distribution of the SNPs and feasibility of parameter estimation critically depends on the network architecture, and in particular the number of hidden nodes in each layer. As frequently seen in applications (Hinton, 2010; Hinton and Salakhutdinov, 2006), we use *p* nodes in the first layer, and *p*/10 nodes in the second layer to achieve a lower-dimensional representation, while still allowing for partitions to be implemented by the approach introduced in the next section.

The number of epochs, i.e. the number of iterations the data is presented to the network, is a tuning parameter which has to be carefully selected in order to avoid overfitting during pre-training and the subsequent refinement. We found that a fixed small number of epochs, such as 20 epochs, worked well in our applications, providing more reliable results, compared to tuning parameter selection based on likelihoods calculated using formula (1), where the analytically intractable partition function *Z*(*θ*) was estimated by annealed importance sampling (Salakhutdinov and Hinton, 2012; Salakhutdinov and Murray, 2008). While such a fixed number of epochs might promote overfitting, we will introduce an approach for strict type 1 error control in subsequent sections, which limits detrimental effects.

Still, we found that useful results were difficult to obtain in situations where the number *p* of visible nodes was larger than a fifth of the number *n* of independent observations. This motivated us to develop the partitioning approach described in the following, which allows to train sub-DBMs on clusters obtained from a partitioning of the SNPs and then reassembling these DBMs into a large DBM.

### 2.2. Partitioning by stagewise regression

Partitioning of restricted Boltzmann machines (RBM) has been suggested for improving training in deep learning (Tosun and Sheppard, 2014). This means that not all parameters of a network are determined at once, but sub-networks are trained for pre-specified partitions of the visible units. To estimate DBMs for a large number of SNPs, we will build on this idea in the following.

The partitioning approach of Tosun and Sheppard (2014) was applied to image input, where partitions could be naturally defined by spatial proximity. For SNP data, spatial proximity could also be considered, but this might focus training on linkage disequilibrium (LD) blocks, i.e. local correlations, which are already rather well known, and lead to overlooking joint patterns of SNPs that are not in close proximity. To prevent this, our partitioning approach is based on a coarse estimate of the joint SNP distribution. The general idea is to coarsely determine multivariable patterns of dependence using a multivariable regression model in a first step, then determine a partition, and subsequently estimate a more accurate DBM model of the joint distribution.

In detail we consider each SNP as a response variable *υ_i_* in a multivariable regression model and want to predict its value by the other SNPs **v**_≠*i*_ that enter the model as covariates, while ignoring interactions and non-linear effects:

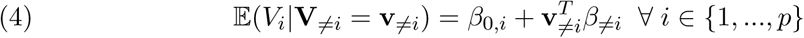

with **v**_≠*i*_ defined as (*υ*_1,…*υ*_*i*−1_,*υ*_*i*+1_,…*υ_p_*_) and *β_≠i_* being the vector of coefficients in the regression model.

Although the models above are deliberately misspecified, having a continuous response form and using artificially standardized binary covariates, one can reasonably expect that strong non-zero relationships can still be identified. Furthermore the advantage of the above mentioned models is that variable selection can be performed in a computationally very efficient way by stagewise regression (Tutz and Binder, 2007) based on covariance statistics. Using stagewise regression we are thus able to select SNPs that are strongly correlated with a given SNP. If *k* steps are performed in this approach, corresponding to the selection of a maximum of *k* covariates with non-zero effects, at most *k* · (*p* — 1) bivariate covariance statistics have to be calculated. This makes computation feasible even for a large number of SNPs, in particular as the covariance statistics can be re-used between the different models.

Stagewise regression, similar to the closely related lasso approach for regularized regression, has the property that it typically assigns only one non-zero estimate to a member of a highly correlated group of covariates that all have an effect (Binder and Schumacher, 2009). In particular for a SNP application, this would mean that only one SNP from an LD block would be selected, and this selection might depend on random variations in the data. For the lasso, the elastic net was introduced to address this (Zou and Hastie, 2005). Specifically, the latter approach exhibits a grouping property, assigning non-zero estimates to each of a correlated group of SNPs that all have an effect. For stagewise regression, an approach for obtaining such a grouping property has been described in Binder and Schumacher (2009). This introduces a second tuning parameter besides the number of steps, which determines the extent to which the grouping property is enforced. Based on the results in Binder and Schumacher (2009), we set this parameter to 0.9 in the following.

Subsequently, we cluster SNPs based on their pairwise relation given by the *p* × *p* matrix *B* of regression coefficients inferred by stagewise regression. Specifically, we construct a distance matrix based on *B*. The matrix *B* is not symmetric and therefore not suitable as a distance matrix, but we can use it to define the symmetric matrix *B*^*^, with 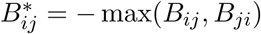 Using *B*^*^ we hierarchically cluster the SNPs using average linkage.

A pre-specified number of clusters is obtained by cutting the resulting tree at an appropriate level, where each cluster represents a SNP group. We suggest to choose the number of clusters such that at least 40 SNPs are clustered together and use this threshold in the following. We train sub-DBMs on each of the SNP groups, resulting in a partitioned DBM, and set the number of the terminal hidden nodes proportional to the number of SNPs in a SNP group. Using a number of *p*/10 second layer hidden units in the joint DBM, this typically allows for partitions with up to *p*/50 groups.

### 2.3. Reassembling an overall DBM

After having obtained parameter estimates for each sub-DBM trained for a group of SNPs in a partition, these DBMs need to be combined into an overall DBM for a joint model of the SNP distribution. In preliminary experiments, we initialized the parameters cross-linking the sub-DBMs by small random values and performed further training iterations. Yet, this did not change performance much, as the parameters from the group-DBMs typically are already larger and dominate subsequent iterations of mean field and Gibbs sampling. Therefore, we suggest to set the cross-linking parameters to zero, i.e. maintaining the partition even in the overall DBM.

### 2.4. Extracting SNP patterns

To extract SNP patterns from a DBM, we consider the association between hidden units and the phenotype of interest in a first step, subsequently identifying visible units, i.e. SNPs, that are potentially associated with hidden units found to be connected to the phenotype. The SNPs found to be potentially associated are themselves tested for association with the phenotype. Both steps looking at the phenotype employ a Bonferroni-Holm (Holm, 1979) correction to maintain type
1 error. Furthermore, to maintain an overall type 1 error of *α*, both steps operate at a level of *α*/2. To test for association between hidden units and a phenotype, we could stochastically determine hidden binary values for each individual from the DBM. Yet, to minimize noise, we deterministically propagate activations in the network to obtain continuous values for each second layer hidden unit. Association with the phenotype is then assessed, by t-tests for binary phenotypes or Cox proportional hazards models for time-to-event phenotypes. To determine SNPs associated with a hidden unit *h*_2*i*_ in the terminal hidden layer **h**_2_ of length *t* that is found to be significantly associated with the phenotype, we again consider multivariable regression, specifically a model

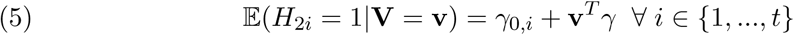

where **v** represents the *p* SNPs. This corresponds to the idea that the hidden unit reflects a multivariable pattern. As in the previous section, the model will certainly be mis-specified, but may nevertheless be useful for variable selection. Accordingly, we perform stagewise regression for estimation as above, using 10 steps to provide a coarse screening. The selected SNPs subsequently are tested for association with the phenotype using the same type of test as for the hidden units.

## 3. Simulation Study

The proposed approach has the potential to identify groups of SNPs that belong to a relevant SNP pattern, and to identify SNPs that might have been missed by standard univariate testing. We anticipate that the performance will depend on the number of SNPs relative to the number of individuals and the complexity of the SNP patterns. Therefore, we designed a simulation study varying these parameters and evaluating performance in terms of SNP identification, with standard univariate testing as a reference.

### 3.1. Design

Binary SNP covariates are generated with a frequency of 0.1 for a value of 1, where the total number of SNPs is 100, 500, 1000, 2000 or 5000. A binary case-control phenotype is generated for a large number of individuals, based on patterns described below, and 500 cases and 500 controls are drawn. We consider two settings with different SNP patterns each involving 50 SNPs, divided into 10 groups containing 5 SNPs each.

In the first pattern (LEVEL1), a case phenotype is generated when in any of the groups at least *k* SNPs have a value equal to 1. Such a pattern might e.g. be seen when there is a large number of biologically relevant genes, where disruption in any of the genes results in a case phenotype, but disruption requires a relatively large number of SNPs. In the second pattern (LEVEL2), the 10 groups are further sub-divided into pairs of groups, each comprising two groups with 10 SNPs in total. A case phenotype is generated when in any of the resulting 5 pairs at least *k* SNPs have a value of 1 in both pair members. Compared to the first pattern, this corresponds to a two level structure with a layered condition on the sum of SNPs. This might e.g. be encountered when disruption of a biological pathway requires disruption of at least two linked genes.

For both the LEVEL1 and LEVEL2 pattern we varied *k*, the number of SNPs per group that were required to be 1 in order to generate a case phenotype, from 3 to 4 and 2 to 3 respectively (called LEVEL1_K3/4, LEVEL2_K2/3 in the following). This results in different average odds ratios across informative SNPs, specifically 1.695 for scenario LEVEL1_K3, 1.896 for LEVEL1_K4, 1.846 for LEVEL2_K2, and 2.336 for LEVEL2_K3. 100 simulation runs were conducted for each simulation setting and each number of simultaneously investigated SNPs.

We performed standard univariate testing using *χ*^2^ tests and Bonferroni correction for an overall level of *α* = 0.05. Using these univariate results as a reference, we investigated the effect of the proposed approach for learning the joint distribution of SNP clusters on the power to detect SNPs that are associated with the phenotype under type 1 error control.

In addition, we considered a combination of the proposed approach and univariate testing, where both are performed at level *α* = 0.025, and results are combined into a joint list, which then satisfies an overall level of *α* = 0.05. This approach reflects the idea that the proposed approach, as a multivariable technique, might be able to extract information that is complementary to the standard univariate approach.

### 3.2. Partitioning performance

The simulated SNP data were partitioned based on stagewise regression and subsequent hierarchical clustering conducted with the estimated regression coefficients *β̂*. In combination with a number of 100 steps, stagewise regression resulted in selection of about 35 (in settings with 100 SNPs) to 80 (in settings with 5000 SNPs) non-zero-effect SNPs. The SNPs were partitioned into on average 1.4 (in settings with 100 SNPs) to 57 (in settings with 5000 SNPs) clusters containing at least 40 SNPs each.

We aimed to learn the joint distribution of binary SNP covariates that were partitioned into different DBMs based on their initially coarsely learned correlation structure. Since in the final DBM all weights between visible and hidden nodes from different clusters are constrained to 0, nothing can be learned about the joint distribution of SNPs whose relation was not detected by the stagewise regression approach. Thus we investigated how well related SNPs are clustered based on the distinctness of the simulated patterns.

To quantify partitioning performance, we determined across how what number of clusters SNPs of a group of related SNPs are distributed. In a perfect clustering scenario all related SNPs would be partitioned into the same DBM cluster. Patterns requiring a larger number of SNPs to be equal to 1, *k* = 4 for LEVEL1 settings and *k* = 3 for LEVEL2 settings, resulted in good partitioning performance (Figure 1). For patterns with a smaller number of SNPs equal to 1, partitioning performed worse, while still at least parts of patterns were recovered in clusters.

**Figure 1.**
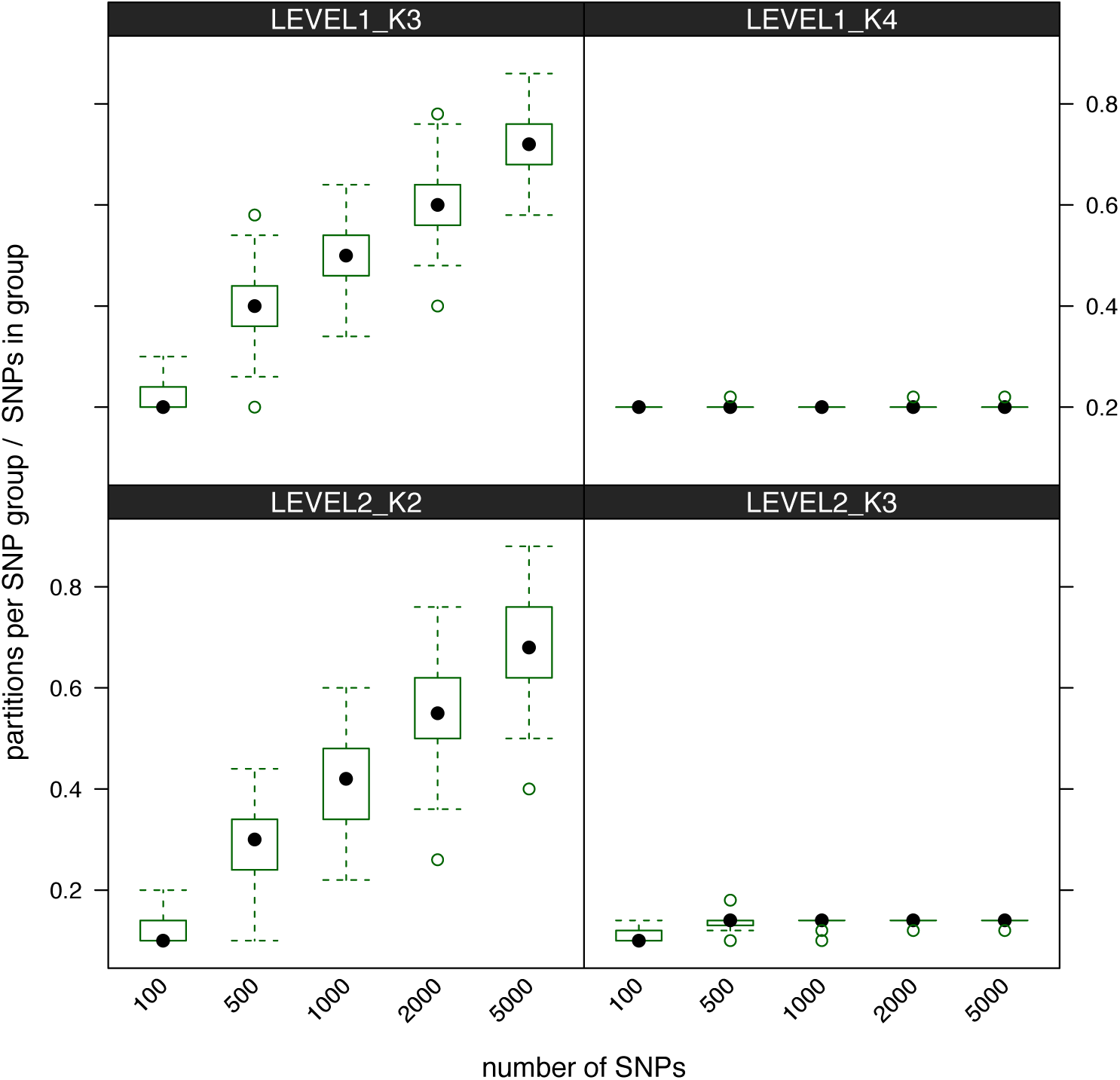
Partitioning performance conditional on the different simulation settings and the number of investigated SNPs. Performance was quantified as the number of partitions in which SNPs of a SNP group were found, divided by the number of SNPs in the simulated SNP group (LEVEL 1 = 5, LEVEL2 = 10). The lower the values are, the better is the partitioning.

### 3.3. SNP identification performance

Using univariate testing as described above, the average number of SNPs that were correctly identified as being associated with the binary phenotype ranged from 1.42 (LEVEL1_K3; 5000 SNPs) to 44.60 (LEVEL2_K3; 100 SNPs) (Figure 2 - “Uni”). As expected these numbers decrease rapidly with the number of SNPs that were investigated and increase with the number of SNPs that were required to generate a case phenotype (lowest for LEVEL1_K3 = 3 SNPs, highest for LEVEL2_K3 = 6 SNPs).

Compared to the univariate analysis, the best performance of the proposed approach is seen for scenarios LEVEL1_K4 and LEVEL2_K3, i.e. when a larger number of SNPs equal to 1 is required for a case pattern (Figure 2 - “partDBM”). While for a small total number of SNPs (*p* = 100) the univariate approach still performs better, the proposed approach is superior for a larger number of SNPs. For the other two scenarios, LEVEL1_K3 and LEVEL2_K2, the univariate approach is superior, albeit at a much smaller number of significant SNPs. This might indicate difficulties of the proposed approach when there is only a weak signal in the data. A further notable property of the proposed approach is that the number of significant SNPs is rather stable irrespective of the total number of SNPs that were investigated, in contrast to univariate testing.

The difficulties of the proposed approach in settings with weak signal (LEVEL1_K3 and LEVEL2_K2) are ameliorated when combining the significant SNPs with those from univariate testing, both obtained at level *α* = 0.025, guaranteeing an overall level of *α* = 0.05. The performance of such a combined approach is equal or superior to univariate testing in almost all scenarios, indicating that the proposed approach can extract complementary information (Figure 2 - “Uni + partDBM”).

**Figure 2.**
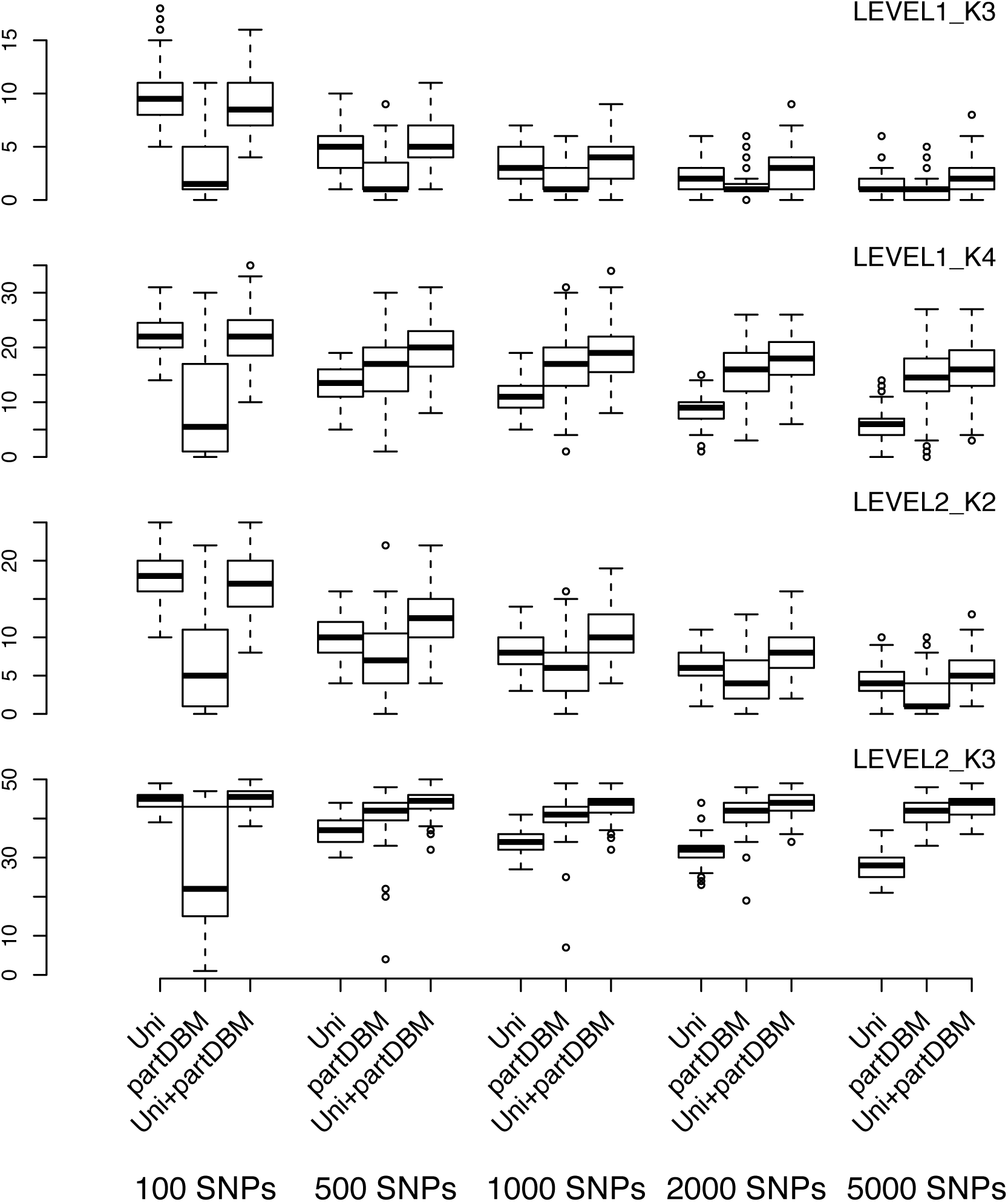
Number of significantly associated SNPs dependending on the number of investigated SNPs and the simulation scenario. Uni = each SNP is tested for association with the case/control phenotype using a *χ*^2^ test, partDBM = visible nodes (SNPs) and the corresponding p-values from the *χ*^2^ test are selected using our partitioned DBM approach, partDBM + Univariate = combination of Uni and partDBM while maintaining the globaf *α* level. P-values were adjusted by the Bonferroni-Holm procedure (FWER ≤ 0.05). 100 simulation runs were conducted for each setting and esch number of SNPs.

## 4. Application

We tested the approach in an application with data from acute myeloid leukemia (AML) patients, where potentially relevant genes were already identified based on gene expression data. The aim was to identify prognostic SNPs, which might provide deeper insight into the underlying biology.

Survival information was available from 308 patients, with 154 deaths and a median survival of 529 days. For each of these patients, 390443 SNPs are available after preprocessing measurements from an Affymetrix SNP 6.0 platform. For more details see Hieke *et al.* (2016a).

As described by Hieke *et al.* (2016b) gene expression measurements are available from a partially overlapping cohort. While in Hieke *et al.* (2016b) the focus had been on identifying gene expression features containing information not already conveyed by the SNP, the present idea is to use the gene expression information to reduce the number of SNPs that are considered for modeling. Specifically, we considered the SNPs mapped to the top seven genes, MAP7, TRIM37, SCAMP4, EXT2, AKT1S1 and MT3, identified by a stagewise regression approach from the gene expression data by Hieke *et al.* (2016b), resulting in a list of 70 SNPs for subsequent modeling by partitioned deep Boltzmann machines.

Partitioning and fitting of DBMs was performed as described above, using four clusters of SNPs, i.e. a partitioning into four sub-DBMs. SNPs were coded into 0/1 values for representing dominant effects. For subsequently identifying relevant hidden units and extracting SNPs, univariate Cox regression models were used, with additive (0/1/2) coding to avoid convergence issues. Significance was assessed with a Wald test.

When fitting Cox models for each of the original 70 SNPs, no SNP was found to be significant after Bonferroni-Holm correction (FWER ≤ 0.05). When considering the top hidden layer of the DBMs, one of the seven hidden units was found to be significantly associated with survival at a level of 0.025 after Bonferroni correction. Sparse regression indicated four SNPs to be associated wth this significant top level hidden unit. After Bonferroni correction at level 0.025, i.e. at an overall level of 0.05, three of these SNPs (rs8082544, rs3826353, rs11656413) were found to be associated with survival. All three SNPs mapped to the gene TRIM37, one upstream and two in the gene body, spanning a total distance of 110508 nucleotides (GRCh38). This spread of location indicates that the proposed approach did not simply identify an LD block, but might have uncovered a more complex pattern.

We validated the identified SNPs using SNP data from AML patients in the Cancer Genome Atlas (TCGA; The Cancer Genome Atlas Research Network (2013); *n* = 200). Each of the SNPs was weakly associated with overall survival (rs8082544: p=0.0235, rs3826353: p=0.0429, rs11656413: p=0.0233; log-rank test). Interestingly, we observed a relation between the cumulative number of at least on risk allele per SNP found in a patient and the patients survival based on Kaplan-Meier estimators (Figure 3) and the results from a Cox regression model (p=0.0289; Wald test). People carrying at least one risk allele of each of the three SNPs had the best survival.

**Figure 3.**
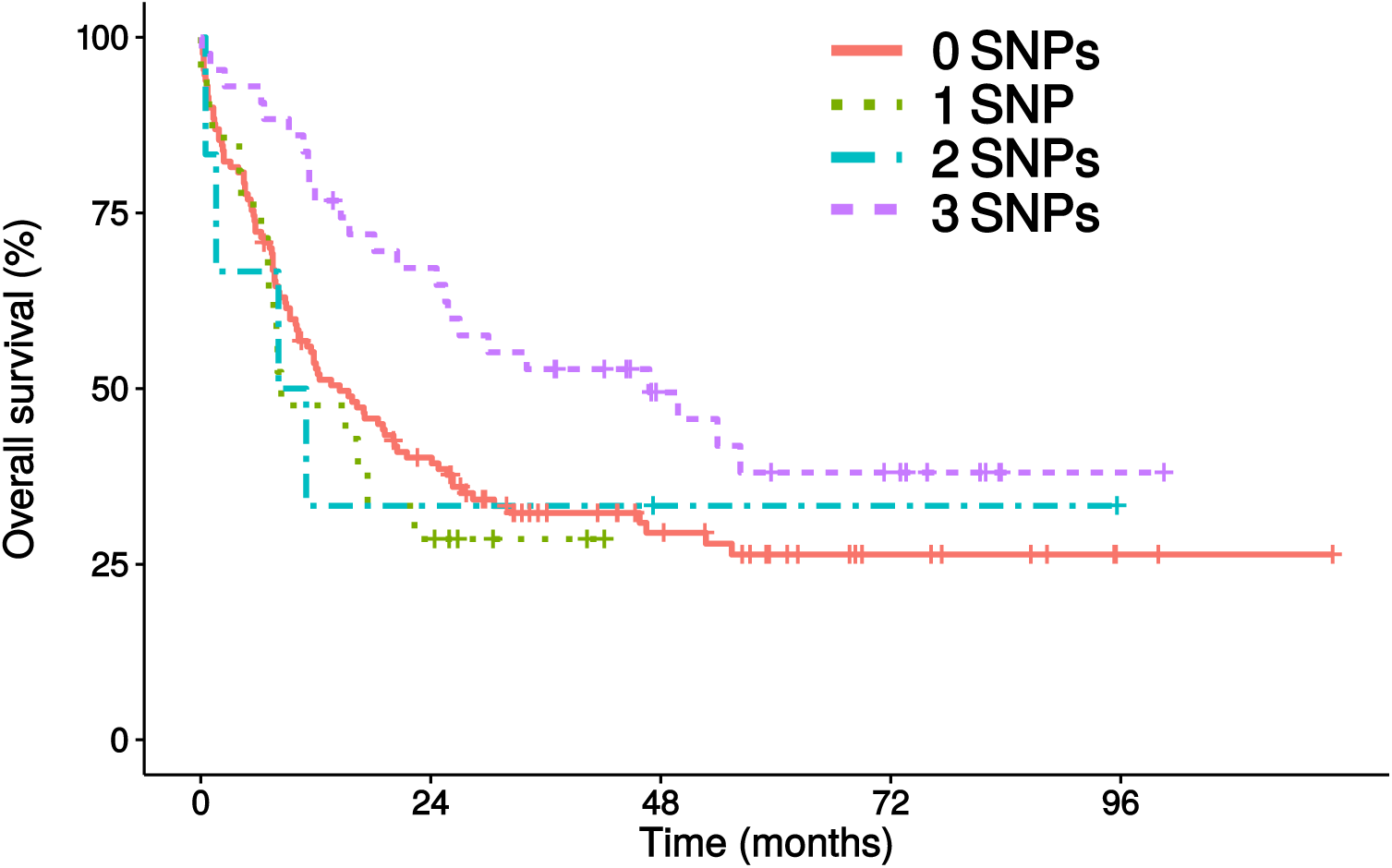
Overall survival in AML patients from independent TCGA validation data (The Cancer Genome Atlas Research Network, 2013). Patients are stratified based on the cumulative number of at least one risk allele of the SNPs rs8082544, rs3826353 and rs11656413 located in TRIM37. Kaplan-Meier estimators are shown.

## 5. Discussion

Although genome-wide association studies (GWAS) have been conducted for more than a decade, joint analysis of multiple SNPs is still not routinely being applied. This indicates the need for approaches to extract low-dimensional features of high-dimensional SNP data based on the underlying distribution that defines SNP co-occurrence. Investigation of haplotype associations (e.g performed in Lambert *et al.*, 2013) might be a valuable approach to reduce dimensionality. However this approach builds on the local correlation structure and consequently is incapable of detecting the joint occurrence of distant risk alleles. Deep learning approaches are a promising technique to progress from local correlation structures to high-level correlation structures but so far have been restricted to applications where the number of observations is often considerably larger than the number of variables. This condition is usually not met in SNP applications.

To apply deep learning techniques in SNP applications, we introduced a partitioning approach that made fitting of deep Boltzmann machines (DBMs) feasible even for a large number of SNPs. Specifically, a sparse regression approach was used for an initial coarse model of the joint SNP distribution, useful enough for obtaining partitions, despite deliberate mis-specification. Extraction of SNPs was subsequently performed by a combination of univariate testing and again sparse regression, to control the type 1 error. The partitioning based on stagewise regression is an important advantage over haploblock-based approaches since in our approach SNPs are jointly investigated based on their co-occurrence and not their physical distance.

In a simulation study we observed that the partitioned DBMs may on average lead to about 1.5 times the number of significant SNPs compared to univariate testing, while also controlling for type 1 error, or almost two times the number of significant SNPs when combining the results from the DBM with the results from univariate testing, indicating that the proposed approach extracts complementary multivariable information. However, we also observed that the performance is strongly dependent on signal strength.

The performance of the initial partitioning was generally worse in the low signal settings. Therefore, in a low signal setting already the initial clustering step might be problematic, meaning that the subsequent training of DBMs cannot be fully successful. Nevertheless, combining the univariate results with the results from the partitioned DBMs did almost never result in worse performance than solely performing univariate testing, indicating that employing partitioned DBMs in addition to univariate testing probably has no adverse effects on power.

We introduced and evaluated our approach for binary SNP values for simplicity, but an extension to 0/1/2 values is rather straightforward. Also we did not consider spurious correlation, such as e.g. found in LD blocks, in the simulation design for evaluation. Yet, preliminary experiments indicated that this would not affect the results much.

In an application to data from AML patients, we considered modeling of SNPs mapped to a set of genes determined to be prognostic based on gene expression data. Such an initial screening step, e.g. based on data from another molecular level, might by a promising approach to reduce the number of SNPs to a level where it can reasonably be modeled by the proposed approach. In our specific application, this allowed for identification of SNPs that would not have been found by standard univariate analyses. These SNPs were validated in a second data set, underlining the robustness of the identified pattern. Interestingly, we observed that the three SNPs showed a cumulative effect on the overall survival, where people carrying at least one risk allele of each of the three SNP loci had the best survival. This strongly supports the validity of our simulation approach which assumed a similar model of multiple SNPs being required to generate a strong phenotype. In addition it highlights that SNPs distributed across a long genomic range can have a strong effect, which might have been missed by haplotype-based approaches.

## 6. Conclusion

Deep architectures are promising techniques for learning compact representations of high dimensional data such as SNP data. Nevertheless the size of current GWAS studies or clinical cohorts, where participant numbers are in the range of thousands or less, would not be sufficient to train a multi-layer DBM for the number of SNPs to be investigated, frequently exceeding the number of studied individuals. To circumvent this issue, we coarsely estimated the relation between SNPs using stagewise regression, and fitted DBMs on resulting small clusters of correlated SNPs. In doing so we effectively constrained the parameter space of the finally merged DBM to ranges that led to learning of meaningful representations. In some settings, these learned representations led to the identification of almost twice the number of significant SNPs when combining results from univariate testing with the significant SNPs identified by the partitioned DBMs. Furthermore in almost no scenario we observed a considerable decrease in the number of significant SNPs compared to solely performing univariate testing. Consequently we think partitioned DBMs are a valuable approach to increase power in genome-wide association studies or clinical cohorts with SNP data. Thus, our partitioning proposal opened the way for adapting a deep learning approach for high-dimensional SNP data. While this application setting certainly requires further investigation, we already anticipate that also other omics settings could potentially benefit from a similar approach.

## Acknowledgements

The results shown here are in part based upon data generated by the TCGA Research Network: http://cancergenome.nih.gov/.

## References

Angermueller, C., Lee, H., Reik, W., and Stegle, O. (2016). Accurate prediction of single-cell dna methylation states using deep learning. BioRxiv, page 055715.

Bengio, Y., Courville, A., and Vincent, P. (2013). Representation learning: A review and new perspectives. IEEE Transactions on Pattern Analysis and Machine Intelligence, 35(8), 1798–1828.

Binder, H. and Schumacher, M. (2009). Incorporating pathway information into boosting estimation of high-dimensional risk prediction models. BMC Bioinformatics, 10(1), 1.

Chen, Y., Lin, Z., Zhao, X., Wang, G., and Gu, Y. (2014). Deep learning-based classification of hyperspectral data. IEEE Journal of Selected Topics in Applied Earth Observations and Remote Sensing, 7(6), 2094–2107.

Chen, Y., Li, Y., Narayan, R., Subramanian, A., and Xie, X. (2016). Gene expression inference with deep learning. Bioinformatics, page btw074.

Ciregan, D., Meier, U., and Schmidhuber, J. (2012). Multi-column deep neural networks for image classification. In 2012 IEEE Conference on Computer Vision and Pattern Recognition (CVPR), pages 3642–3649. IEEE.

Graves, A., Mohamed, A.-r., and Hinton, G. (2013). Speech recognition with deep recurrent neural networks. In 2013 IEEE International Conference on Acoustics, Speech and Signal Processing, pages 6645–6649. IEEE.

Hieke, S., Benner, A., Schlenk, R. F., Schumacher, M., Bullinger, L., and Binder, H. (2016a). Identifying prognostic snps in clinical cohorts: Complementing univariate analyses by resampling and multivariable modeling. PloS One, 11(5), e0155226.

Hieke, S., Benner, A., Schlenl, R. F., Schumacher, M., Bullinger, L., and Binder, H. (2016b). Integrating multiple molecular sources into a clinical risk prediction signature by extracting complementary information. BMC Bioinformatics, 17(1), 327.

Hinton, G. (2010). A practical guide to training restricted boltzmann machines. Momentum, 9(1), 926.

Hinton, G. E. (2002). Training products of experts by minimizing contrastive divergence. Neural Computation, 14(8), 1771–1800.

Hinton, G. E. and Salakhutdinov, R. R. (2006). Reducing the dimensionality of data with neural networks. Science, 313(5786), 504–507.

Holm, S. (1979). A simple sequentially rejective multiple test procedure. Scandinavian Journal of Statistics, pages 65–70.

Krizhevsky, A., Sutskever, I., and Hinton, G. E. (2012). Imagenet classification with deep convolutional neural networks. In Advances in Neural Information Processing Systems, pages 1097–1105.

Lambert, J.-C., Grenier-Boley, B., Harold, D., Zelenika, D., Chouraki, V., Kamatani, Y., Sleegers, K., Ikram, M. A., Hiltunen, M., Reitz, C., et al. (2013). Genome-wide haplotype association study identifies the frmd4a gene as a risk locus for alzheimer’s disease. Molecular Psychiatry, 18(4), 461–470.

Leung, M. K., Xiong, H. Y., Lee, L. J., and Frey, B. J. (2014). Deep learning of the tissue-regulated splicing code. Bioinformatics, 30(12), i121–i129.

Neal, R. M. and Hinton, G. E. (1998). A view of the em algorithm that justifies incremental, sparse, and other variants. In Learning in Graphical Models, pages 355–368. Springer.

Quang, D., Chen, Y., and Xie, X. (2014). Dann: a deep learning approach for annotating the pathogenicity of genetic variants. Bioinformatics, page btu703.

Salakhutdinov, R. and Hinton, G. (2012). An efficient learning procedure for deep boltzmann machines. Neural Computation, 24(8), 1967–2006.

Salakhutdinov, R. and Hinton, G. E. (2009). Deep boltzmann machines. In AISTATS, volume 1, page 3.

Salakhutdinov, R. and Murray, I. (2008). On the quantitative analysis of deep belief networks. In Proceedings of the 25th International Conference on Machine Learning, pages 872–879. ACM.

The Cancer Genome Atlas Research Network (2013). Genomic and epigenomic landscapes of adult de novo acute myeloid leukemia. New England Journal of Medicine, 368(22), 2059–2074.

Tosun, H. and Sheppard, J. W. (2014). Training restricted boltzmann machines with overlapping partitions. In Joint European Conference on Machine Learning and Knowledge Discovery in Databases, pages 195–208. Springer.

Tutz, G. and Binder, H. (2007). Boosting ridge regression. Computational Statistics & Data Analysis, 51(12), 6044–6059.

Wei, W.-H., Hemani, G., and Haley, C. S. (2014). Detecting epistasis in human complex traits. Nature Reviews Genetics, 15(11), 722–733.

Zou, H. and Hastie, T. (2005). Regularization and variable selection via the elastic net. Journal of the Royal Statistical Society: Series B (Statistical Methodology), 67(2), 301–320.

